# Expanding the iModulon Knowledgebase for *Bacillus subtilis*: Updated Transcriptome Decomposition Reveals Novel Regulatory Mechanisms

**DOI:** 10.1101/2025.09.29.679290

**Authors:** Peter Gockel, Ivan Mijakovic, Bernhard Palsson, Daniel Zielinski, Emre Özdemir, Lei Yang

## Abstract

The Gram-positive bacterium *Bacillus subtilis* is widely used as a model organism for studying cell differentiation, microbial physiology and gene regulation. Its genetic accessibility and strong protein secretion machinery make it an excellent host for industrial protein production. Transcriptomics has become an essential tool in understanding these traits and the organism’s response to environmental stimuli. Recent advantages in transcriptomics have enabled genome-wide analysis of gene expression patterns under various environmental conditions and genetic perturbations, interpreting such data, however, remains challenging. Independent Component Analysis (ICA) has emerged as a powerful method to decompose bacterial transcriptomes into sets of co-regulated genes – so-called iModulons. In this study, we present an updated iModulon decomposition of *Bacillus subtilis* transcriptomes, generated from a compendium of 782 high-quality RNAseq samples. This dataset includes a wide range of environmental and genetic conditions. ICA decomposition revealed 142 iModulons, that captured 80% of the expression variance. We compare our version of the modularized *B. subtilis* transcriptome with previous decompositions and show that the updated composition enhances our understanding of *B. subtilis* gene regulation and enables novel insights into the organism’s regulon structure and gene memberships of the WalR regulon.

## 2. Introduction

The transcriptional regulatory network (TRN) represents a crucial mechanism inside cells, that enables the controlled expression of genes in response to environmental conditions like the availability of carbon sources, changes in temperature or the presence of antibiotics, as well as genetic perturbations. Within a TRN, the expression of genes is regulated by the binding of transcription factors (TFs) on regulatory motifs on the DNA (Saint-André, 2021). Groups of genes that are co-regulated by the same transcription factor are often considered to be part of the same regulon that is activated or repressed in response to environmental stimuli. Studying TRN structures in bacteria enables the mechanistic understanding of larger cellular processes like cellular differentiation or stress responses to unfavorable environmental conditions.

*Bacillus subtilis*, a Gram-positive bacterium, is considered a model organism for studying such processes. The organism has been instrumental in advancing our understanding of bacterial physiology, gene expression and regulation, and cellular differentiation (Stülke et al., 2023), (Harwood, 1992). In recent decades, the organism utility has extended into biotechnology, where it is used as a microbial cell factory, among other examples, most notably to produce industrial enzymes and other proteins (Liu & Yu, 2025), (Westers et al., 2004). The organism presents an excellent host for these types of applications due to its capacity to efficiently secret proteins and its generally regarded as safe (GRAS) status (Gu et al., 2018). A large part of our understanding of *B. subtilis* physiology and gene regulation can be attributed to the application of high-throughput transcriptomics workflows like the generation of microarray or RNAseq datasets.

The increasing availability of these datasets has opened new possibilities for dissecting transcriptional regulations on a systems level with RNAseq enabling precise quantification of gene expression in bacteria like *B. subtilis* in response to changing environmental or genetic conditions. Interpreting these large datasets, however, remains a major challenge, especially when trying to uncover latent regulatory structures and functional coherent gene modules.

One approach that has proven valuable in addressing this challenge is Independent Component Analysis (ICA), an unsupervised machine learning approach, that decomposes transcriptomics datasets into sets of independently regulated sets of genes – so-called iModulons – whose expression patterns vary together across conditions (Sastry et al., 2019). iModulons often represent regulons, stress responses or metabolic modules. They provide an interpretable and compact view of an organism transcriptional regulatory network (TRN). ICA has been applied to capture the iModulon composition of multiple bacterial species and made available on iModulonDB (Catoiu et al., 2025), including *Escherichia coli*, *Pseudomonas putida*, *Vibrio natriegens* and *Streptomyces albidoflavus (Sastry et al., 2019), (Shin et al., 2023), (Jönsson et al., 2025)*.

Rychel et al. (2020) applied ICA to a high-quality *B. subtilis* microarray dataset, identifying 83 iModulons that captured a major part of the transcriptomic variance and aligned closely with known regulons. This study demonstrated that ICA could uncover biologically meaningful, independently modulated gene sets and offered new insights into condition-dependent regulatory mechanism – for instance, revealing a string activation of the tryptophan synthesis iModulon under ethanol stress, suggesting a previously unrecognized link between ethanol exposure and tryptophan depletion in *Bacillus subtilis* (Rychel et al., 2020).

Building on this, Sastry et al. (2024) developed the iModulonMiner pipeline and PyModulon toolkit and used them to exemplary reconstruct the *B. subtilis* iModulon structure from an expanded RNAseq compendium mined from publicly available NCBI SRA transcriptomes up until the year 2020 (Sastry et al., 2024). This study revealed 72 robust iModulons that together explain 67% of the total expression variance. Of these, 52 iModulons were enriched for known transcriptional regulators while the rest included functional modules, single-gene artifacts, or uncharacterized gene sets. This iModulon structure provides a data-driven view of the *B. subtilis* transcriptome Yet, as with every data-driven model, the quality and scope of the resulting iModulons are inherently tied to the breadth and diversity of the underlying dataset. In recent years an increasing number of publicly available RNAseq datasets for *B. subtilis* have become available, representing a broader range of experimental conditions, stress responses, nutrient limitations, genetic perturbations, and industrially relevant growth environments. These new datasets provide an opportunity to revisit the existing iModulon structure with greater statistical power and biological diversity. Incorporating such data not only refines existing iModulons but also enables the discovery of previously undetected regulatory modules or the assignment of new gene members to known regulons.

In this study, we updated the iModulon structure of *Bacillus subtilis* by applying ICA to an expanded, curated RNAseq compendium comprising 782 high-quality samples. This analysis resulted in 142 robust iModulons, which together explain 80% of the variance in gene expression. We compare the updated model to the previous published decomposition by Sastry et al (Sastry et al., 2024), examining overlap in gene content, associated regulators, and functional enrichment. The updated structure preserved many known regulator-associated iModulons while revealing 53 novel iModulons, including several active under conditions that were underrepresented in earlier datasets. To showcase how a data-driven approach to representing TRN of *B. subtills* and other organism can help refine what is previously known about regulatory relationships, we looked closer into the case of the WalR iModulon. The WalR iModulon captures, besides previously known regulon gene members, the genes *slp* and *ykrP*, previously not known to be regulated by WalR. A closer look into the gene’s regulatory regions, confirmed the presence of potential WalR regulator DNA binding motifs. Experimental validation revealed that mutagenesis of the proposed regulatory sites, alleviates the WalR-mediated activation of these genes.

## 3. Materials & Methods

### 3.1 Acquisition and processing of RNAseq data from NCBI SRA

The RNAseq samples using in this study were obtained from NCBI SRA. Initially a total of 2340 transcriptomes samples from Bacillus subtilis were mined from SRA using the workflow described by Sastry et al (Sastry et al., 2024), which is available at https://github.com/SBRG/iModulonMiner. In short, metadata for the 2340 RNAseq samples was downloaded from NCBI SRA using the query “Bacillus subtilis”. Next, reads were trimmed using Trim galore (https://www.bioinformatics.babraham.ac.uk/projects/trim_galore/) with default settings. Following this FastQC (https://www.bioinformatics.babraham.ac.uk/projects/fastqc/) was applied, followed by aligning the reads to the *Bacillus subtilis* 168 reference genome from NCBI (AL009126.3) using Bowtie (Langmead & Salzberg, 2012). After this, the direction of reads was inferred using RSEQC (Wang et al., 2012), read counts were generated by applying featureCounts (Liao et al., 2014) and all quality control metrics were compiled using MultiQC (Ewels et al., 2016).

### 3.2 Quality Control and data normalization

Next, all compiled samples were quality controlled based on their FastQC metrics. Samples that failed one of the following quality criteria were discarded before further analysis: per base sequence quality, per sequence quality score, per base n content, and adapter content. Additionally, all samples that had less than 500.000 reads mapped to their containing coding sequences were removed. Furthermore, hierarchical clustering was used to remove samples that don’t conform to the global expression profiles. Next, we used manual curation to ensure proper metadata availability for the remaining samples. All samples that could not be linked to sufficient information regarding there precise sampling conditions, were discarded. Lasty, only samples with at least two replicates and a Pearson R correlation of > 0.95 between these replicates were further analyzed or otherwise discarded. The final compendium comprised of 782 high-quality samples that were normalized on a per-project bases, during which a reference condition was defined for each of the projects the respective samples originated from. This step was included to ensure obviation of batch effects and enables that independent components are found due to biological rather than technical variation.

### 3.3 Independent Component Analysis and iModulon Characterization

To obtain independent regulated sets of genes from the expression dataset, we applied the independent component analysis algorithm optICA, an extension of the widely used FastICA algorithm (Hyvarinen, 1999), to find robust components at the optimal dimensionality (McConn et al., 2021). Dimensions were selected from 60 to 320 with a step size of 20 and ICA was run sequentially. The optimal number of dimensions is selected as the dimensionality in which the number of single-gene iModulons is the same a s the number of final components (figure S1). The optimal dimensionality used in the study was found to be 280, resulting in 142 robust components. During ICA the gene expression matrix X gets decomposed into a gene weight matric M and the iModulon activity matrix A (X=M*A). The M matric contains the weights of genes above the gene weight threshold for each iModulon. The A matrix represents the activity of each of those iModulons over the condition space.

Once the optimal components were obtained, the resulting iModulons were characterized using the PyModulon package (Sastry et al., 2024). Since a well annotated TRN is available from SubtiWiki (Elfmann et al., 2025), known TRN information was used for iModulon enrichment, following the steps described in the iModulonMiner workflow (https://github.com/SBRG/iModulonMiner/tree/main/5_characterize_iModulons)

### 3.4 Strains, plasmids, oligonucleotides, media and growth conditions

All bacterial strains, plasmids, BioBricks and oligonucleotides used in this study are listed in the tables. *E. coli* and *B. subtilis* cells were grown in Luria-Bertain (LB) medium. Antibiotics were used at the following concentrations: ampicillin at 100 µg/ml; chloramphenicol at 5 µg/ml.

**Table 1:** Bacterial strains used in this study.

| Strain | Parental | Relevant genotype | Reference, source |
| --- | --- | --- | --- |
| 1A1 | N/A | Strain 168, trpC2 | BGSC |
| Bs118 | 1A1 | + pGO127 | This study |
| Bs119 | 1A1 | + pGO128 | This study |
| Bs120 | 1A1 | + pGO129 | This study |
| Bs121 | 1A1 | + pGO130 | This study |
| Bs122 | 1A1 | + pGO131 | This study |
| Bs123 | 1A1 | + pGO132 | This study |

**Table 2:** Plasmids used in this study.

| Plasmid | Parental | Description | Reference/ source |
| --- | --- | --- | --- |
| pDL | N/A | *amyE* integration plasmid for transcriptional fusions to *bgaB* | (Yuan & Wong, 1995)BGSC |
| pGO127 | pDL + BB286 | pDL plasmid containing the regulatory region upstream of *slp* fused to *bgaB* | This study |
| pGO128 | pDL + BB289 | pDL plasmid containing the regulatory region upstream of *ykrP* fused to *bgaB* | This study |
| pGO129 | pDL + BB294 | pDL plasmid containing the mutated (ACA to TGT) regulatory region upstream of *slp* fused to *bgaB* | This study |
| pGO130 | pDL + BB295 | pDL plasmid containing the mutated BM-1 (ACA to TGT) regulatory region upstream of *ykrP* fused to *bgaB* | This study |
| pGO131 | pDL + BB296 | pDL plasmid containing the mutated BM-2 (ACA to TGT) regulatory region upstream of *ykrP* fused to *bgaB* | This study |
| pGO132 | pDL + BB298 | pDL plasmid containing the mutated BM-1 and BM-2 (ACA to TGT, respectively) regulatory region upstream of *ykrP* fused to *bgaB* | This study |

**Table 3:** BioBricks constructed in this study. Bs gDNA = genomic DNA extracted from the Bacillus subtilis 1A1 strain.

| BioBrick ID | Template | Oligonucleotides | Reference |
| --- | --- | --- | --- |
| BB286 | Bs gDNA | PGO375, PGO376 | This study |
| BB287 | Bs gDNA | PGO375, PGO277 | This study |
| BB288 | Bs gDNA | PGO378, PGO376 | This study |
| BB289 | Bs gDNA | PGO379, PGO380 | This study |
| BB290 | Bs gDNA | PGO379, PGO383 | This study |
| BB291 | Bs gDNA | PGO379, PGO381 | This study |
| BB292 | Bs gDNA | PGO382, PGO380 | This study |
| BB293 | Bs gDNA | PGO384, PGO380 | This study |
| BB294 | BB287, BB288 | PGO375, PGO376 | This study |
| BB295 | BB291, BB292 | PGO379, PGO380 | This study |
| BB296 | BB290, BB293 | PGO379, PGO380 | This study |
| BB297 | BB295 | PGO379, PGO383 | This study |
| BB298 | BB297, BB293 | PGO379, PGO380 | This study |

**Table 4:**
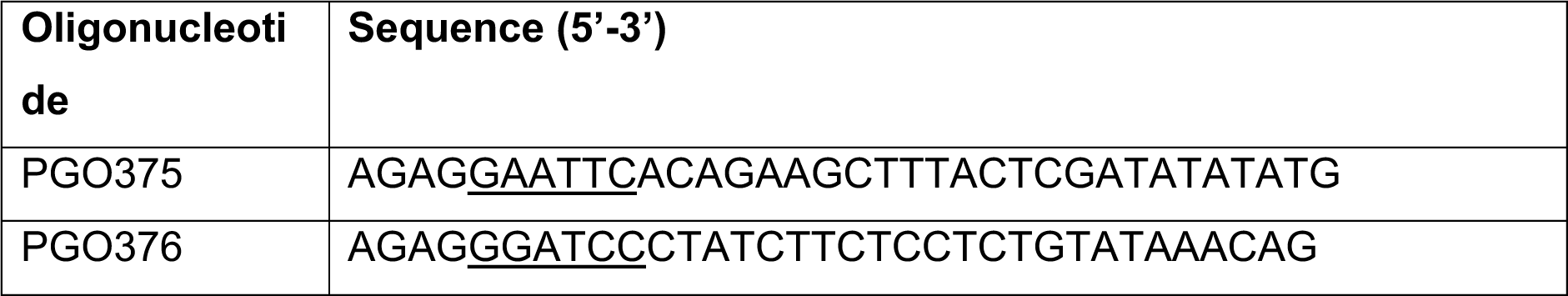
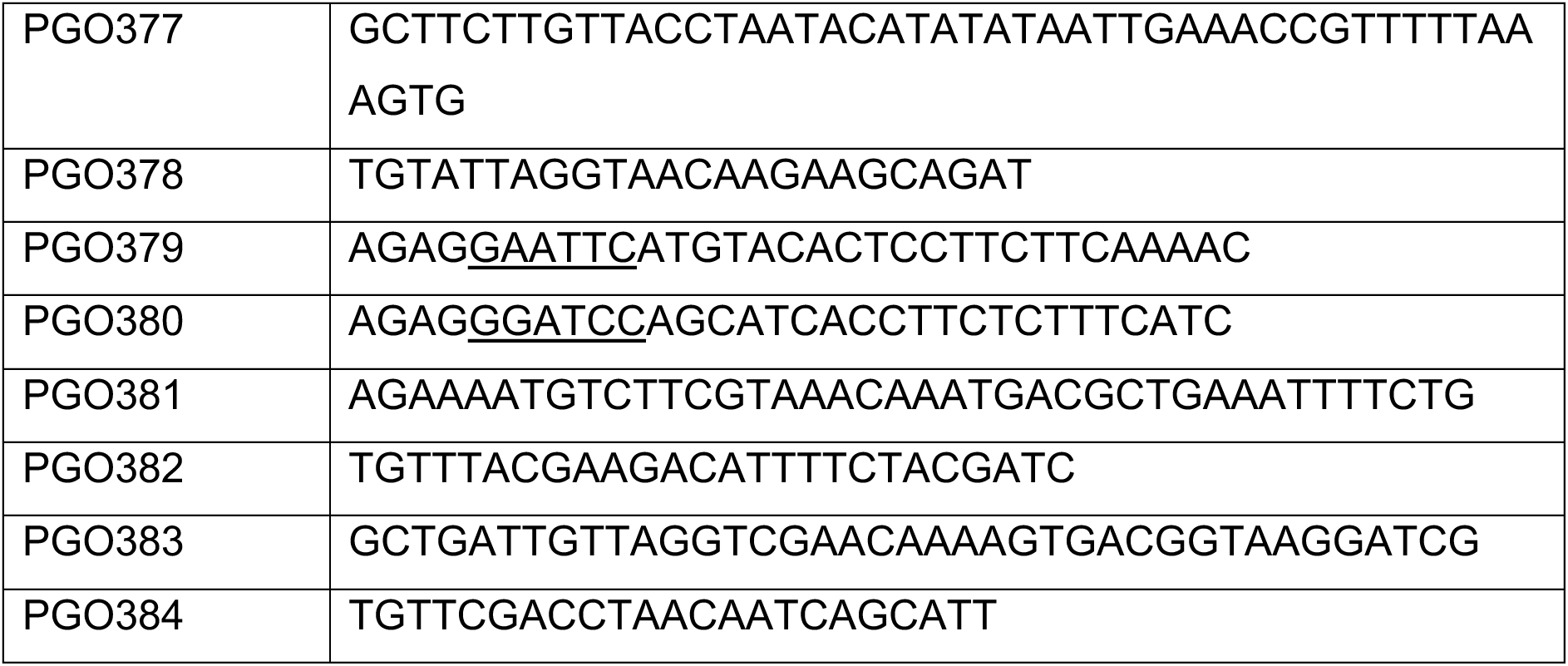
Oligonucleotides used in this study. The restriction sites introduced are indictated as underlined nucleotides.

### 3.5 Construction of plasmids with bgaB fusion to regulatory region of genes of interest

The pDL-based plasmids pGO127, pGO128, pGO129, pGO130, pGO131, pGO132 and pGO133 for the fusion of the upstream regulatory regions of *slp* and *ykrP* to the thermostable e gene *bgaB* were constructed by amplifying the respective regulatory regions from genomic DNA of Bacillus subtilis strain 1A1. The construction of the plasmids was performed like previous reported (Huang et al., 2013). In short, the indicated primers were used to attached EcoR1 and BamH1 restriction enzyme recognition sites to the amplified genomic regions. The resulting fragments and the pDL backbone were digested with EcoR1 and BamH1 and the regulatory regions cloned into the vector. In order to introduce mutations into the binding motifs of the regulatory regions, the regions were amplified in two separated fragments using primers introducing the mutation into the region of interested and generated a 20 bp overlap between the two fragments. Using overlap extension PCR the two fragments were fused and the successful introduction of the mutation verified using Sanger sequencing upon plasmid cloning.

### 3.6 Transformation of *Bacillus subtilis*

For the transformation of *B. subtilis* strain 1A1 with plasmid DNA, a protocol previously described was used. In short, a single colony from the freshly streaked strain of LB-Agar used to inoculate 10 ml of SM1 medium (2 g/L ammonium sulphate, 14 g/L dipotassium hydrogen phosphate, 6 g/L potassium dihydrogen phosphate, 0.7 g/L sodium citrate, 0.2 g/L magnesium sulfate heptahydrate, 2 g/L yeast extract, 0.25 g/L casamino acids and 5 g/L glucose) in a 50 ml falcon tube. The culture was incubated at 37 °C and 250 rpm shaking for 16 hours and subsequently used to inoculate 10 ml of SM1 medium in a 250 ml shake flask to a starting OD of 0.5. The culture was incubated at 37 °C and 250 rpm for 3 hours. Next, 10 ml of SM2 medium (2 g/L ammonium sulphate, 14 g/L dipotassium hydrogen phosphate, 6 g/L potassium dihydrogen phosphate, 0.7 g/L sodium citrate, 0.8 g/L magnesium sulfate heptahydrate, 1 g/L yeast extract, 0.1 g/L casamino acids, 5 g/L glucose, 2.25 mM CaCl_2_) were added to the culture and the culture was incubated for another 2 hours under the same cultivation conditions as previously described. Next, the culture was distributed into 500 µl aliquots in 2 ml reaction tubes and 250 ng of plasmid DNA was added. The mixture was incubated for 30 mins at 37 °C with 800 rpm shaking in a thermomixer before 300 µl of LB medium was added to allow the cells to recover. After 2 h of incubation, the cells were plated on LB agar supplemented with 5 µg/ml of chloramphenicol and incubated at 37 °C over night. On the next day, successful integration of the desired construct into the amyE locus was confirmed by colony PCR.

### 3.7 Activity assay of thermostable β-galactosidase

The activity of the thermostable β-galactosidase BgaB was measured as previously described (Huang et al., 2013). In short, *B. subtilis* strains were grown over night in 10 ml LB medium supplemented with 5 µg/ml of chloramphenicol (LB + cmp) at 37 °C and 250 rpm shaking. The next day, 40 ml of fresh LB + cmp in a 250 ml shake flask was inoculated to an OD of 0.1 using the overnight culture. Once the culture reached an OD of 0.4 the cells were split into two fractions of 20 ml each in two separate 250 ml shake flasks (timepoint 0). One fraction was incubated at 37 °C and the second fraction at 51 °C with 250 rpm shaking respectively. The cells grown for various times as indicated in figure 5a&b. At each timepoint a sample of 2 ml of bacterial culture was pelleted and resuspended in an equivalent volume of BgaB buffer (25 mM potassium phosphate at pH 6.4, 50 mM KCl, and 1 mM MgSO_4_). 1 ml of the sample was transferred into a curvette for OD measurement at 600 nm. The remaining 1 ml was used for the BgaB assay. Cells were permeabilized by addition of 50 µl 1% sodium dodecyl sulfate and 100 µl of chloroform, followed by vortexing. Subsequently, the reaction was initiated by the addition of 0.2 ml o-nitrophenol-β-D-galactopyranoside at a concentration of 4 mg/ml. The reaction mixture was incubated at 55 °C for 30 mins and stopped by the addition of 0.5 ml Na_2_CO_3_. Following centrifugation to remove chloroform and cell debris, 200 µl of the supernatant were transferred into a 96-well plate to measure the absorbance at 420 nm and 55 nm. BgaB activity is represented in Miller units.

## 4. Results

### 4.1 Construction of an expanded *B. subtilis* expression compendium from publicly available NCBI SRA data

To update the previous decomposition of *B. subtilis* transcriptomes we collected publicly available transcriptome data from the National Center for Biotechnology Information (NCBI) Sequence Read Achieve (SRA) (Katz et al., 2022) and processed them according to the pipeline described previously (Sastry et al., 2024). In short, this pipeline consists of downloading publicly available RNAseq samples from NCBI SRA, processing these sequencing reads, filtering them based on strict quality standards, applying the optICA algorithm and lastly characterizing the resulting independent components (Figure 1a).

**Figure 1:**
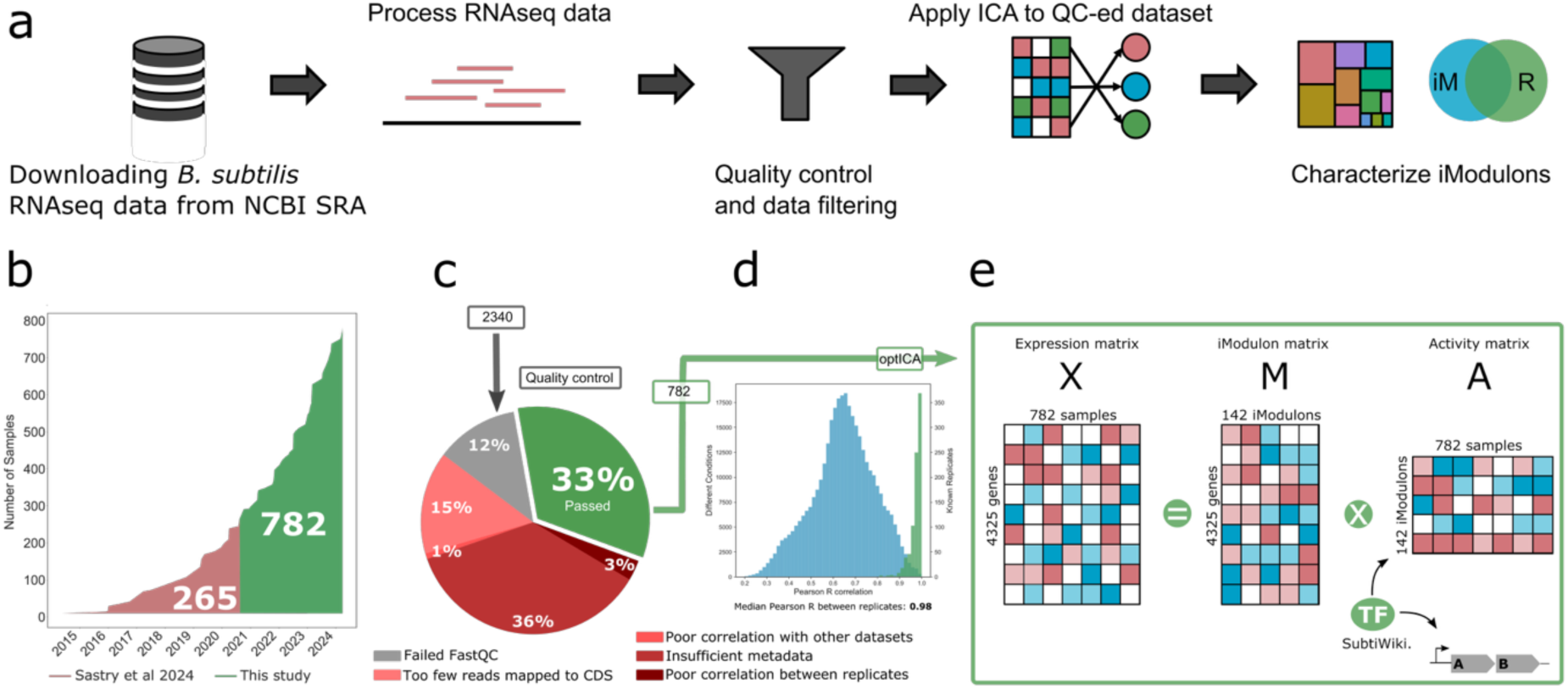
Overview of workflow and RNAseq samples used to update the iModulon decomposition of Bacillus subtilis. (a) Graphical representation of iModulonMiner workflow applying ICA to quality-controlled dataset of RNAseq samples ubtained from NCBI SRA. (b) Overview of high-quality (after QC) RNAseq samples deposited in NCBI SRA and datasets integrated in the previous iModulon decomposition by Sastry et. al (red) and newly added samples in this study (green). (b) Overview of quality control steps during samples processing, which yielded a total of 782 high-quality samples curated from 2340 input samples. (d) Median Pearson R between replicates in the 782-sample-dataset of 0.98. (e Schematic representation of iModuon decomposition of expression patterns of 4325 *Bacillus subtilis* genes over 782 samples (X matrix), decomposed into 142 robust iModulons represening the 4325 *B. subtilis* genes (M matrix) and there associated activities (A matrix).

Initially a total of 2340 RNAseq samples were obtained from the SRA database and were subjected to data processing and quality control steps. During this procedure approximately two thirds of the samples (1558 samples) were excluded after failing to meet one of the following criteria: FastQC quality criteria, insufficient reads mapped to the coding sequences (> 500.000), poor global correlation, insufficient metadata availability and poor correlation between biological replicates. Out of these, the insufficient availability of metadata, accurately describing information on cultivation conditions during RNA sampling, was the largest cause of dataset exclusion (Figure 1c). Upon sample filtering, a curated high-quality compendium of 782 RNAseq datasets was used for further analysis. While previous iModulon decompositions of *Bacillus subtilis* transcriptomes incorporated 265 samples, this updated compendium presented an almost 3-fold increase in input data for the subsequent ICA-decomposition which can be largely attributed to the large increase in publicly available datasets in recent years (Figure 1b) (Sastry et al., 2024).

The curated set of 782 RNAseq samples showed a high median Pearson R of 0.98 between replicates (Figure 1d) and represented a wide range of environmental conditions, such as supplementation of various carbon sources or antibiotics, cultivation temperatures and cell differentiation stages; as well as genetic perturbations such as, knockouts of transcription factors. In the next step, the optICA algorithm was applied to find robust independent components at optional dimensionality. Here, the 4325 genes of expression pattern *Bacillus subtilis* across the 782 samples (**X** matrix) were decomposed into 142 iModulons (**M** matrix) and their activity profile across the samples (**A** matrix) (figure 1e). TRN information and gene annotations from SubtiWiki (Elfmann et al., 2025) were applied to characterize the resulting iModulons, which were subsequently annotated based on their overlap with known regulons.

The increased diversity of the conditions in this extended dataset, compared to previous iModulon decompositions, ensures that the resulting iModulons represent both the core physiological states of *B. subtilis* and specialized responses that were previously underrepresented. Compared to earlier efforts, such as the microarray decomposition (Rychel et al., 2020) or the previous RNAseq decomposition, the curated dataset here not only more than doubles the sample size but also incorporates new experimental designs and stress regimes, broadening the observable regulatory landscape.

### 4.2 Independent component analysis reveals 142 iModulons describing the TRN of *Bacillus subtilis*

Applying the optICA algorithm to this dataset resulted in a set of 142 robust iModulons that described the transcriptional regulatory network of *Bacillus subtilis* through a top down approach and its response to various environmental conditions including growth, supplementations, temperatures, knockouts or media compositions. The newly obtained number of iModulons presents a nearly 2-fold increase in number of iModulons compared to the previous decomposition constructed by Sastry et. al. To characterize the resulting iModulons in respect to their biological function within the *Bacillus subtilis* regulatory network, we enriched the signals with TRN information from SubtiWiki (Elfmann et al., 2025), resulting in 114 regulatory iModulon that describe known regulatory functions in the organism, 18 uncharacterized iModulons and 5 single-gene iModulons. Furthermore, SubtiWiki functional categories were used to group the iModulons into functional group based on their enriched regulator and the known function of the genes contained in the respective iModulon. Out of these the largest group represents iModulons associated to amino acid and nucleotide metabolism, while carbon and misc. metabolism-related iModulons present the two second largest category groups observed. A full overview of all iModulons captured from this updated RNAseq compendium can be seen in the treemap in figure 2a.

**Figure 2:**
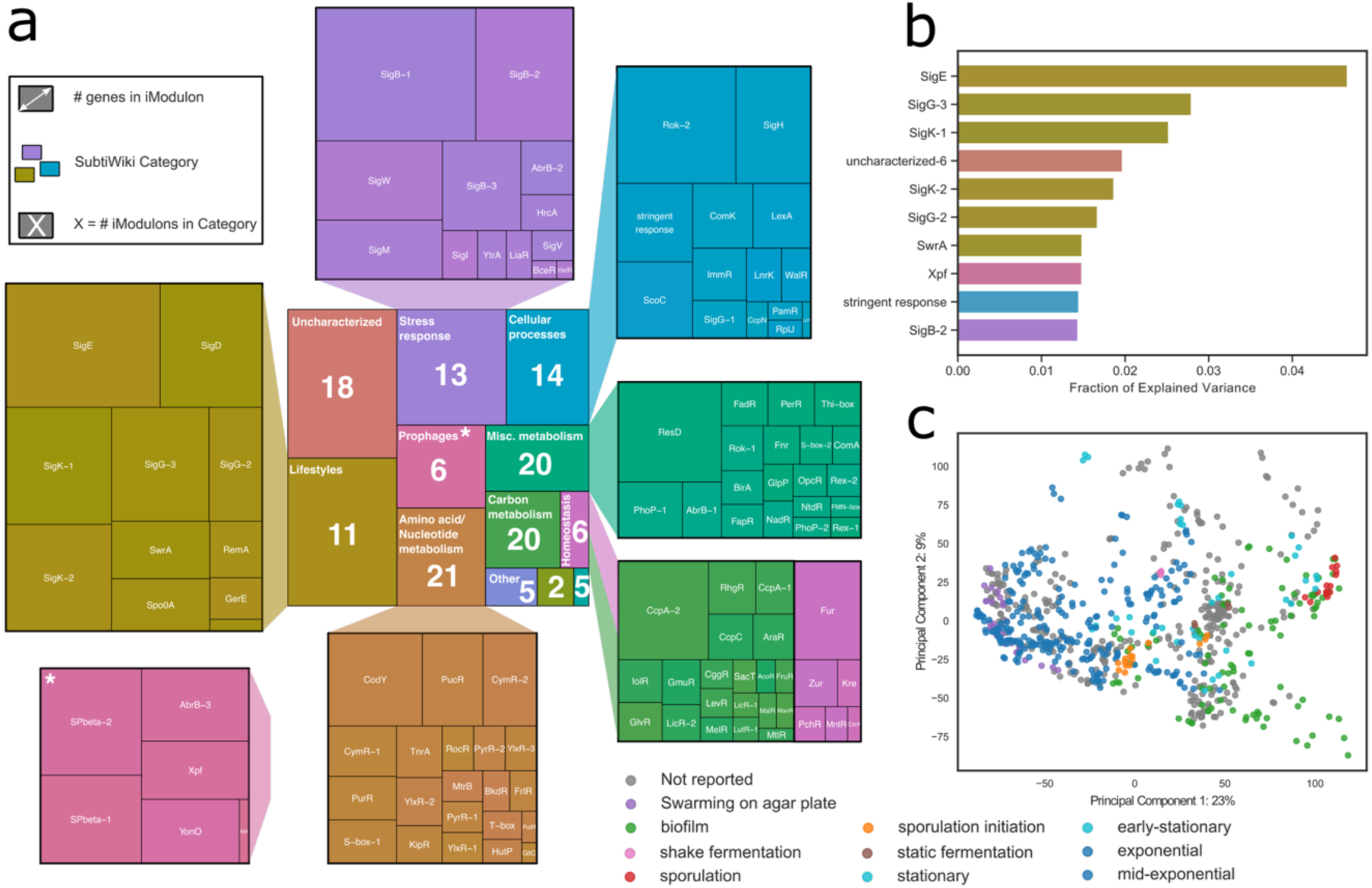
Overview of the capture 142 iModulons enriched with known TRN information. (a) Treemap of the iModulon decomposition, in which each color represents a SubtiWiki category that the respective iModulons could be assciated with. The size of each rectangle represents the number of genes captured in each iModulon. The numbers inside the center rectangles represented the number of iModulons in the respective category. iModulons were enriched with known TRN informaton from SubtiWiki. (b) Top 10 iModulons explaining the largest portions of the expression variance in the underlying dataset. (c) Scatter plot of the two largest components from Principal Component Analysis (PCA), in whicb samples are colored by their growth phase parsed from literature.

SigB-1 was the largest captured iModulon while SigE iModulon explained the largest portion of the expression variance in the dataset (Figure 2a, 2b). Many of the iModulons found among the top 10 iModulons explaining the largest portion of the expression variance are associated to the lifestyle SubtiWiki category, presumable due to the large number of experimental conditions exploring transcriptional responses to cellular differentiation processes and cellular growth phases like sporulation or biofilm formation within the input dataset. A full overview of the contribution of each iModulon to the explain variance can be found in supplementary data (Figure S2).

Principle component analysis (PCA) further supports this by showing that the two largest components were able to separate the samples based on the differential iModulon activity in the samples because of the growth phase the cells were in when sampling. This can be seen in Figure 2c in which samples are colored based on their growth phase found in the literature of the associated studies.

### 4.3 Mapping iModulons to their known corresponding regulons

To evaluate the effectiveness of this approach, we examined how well the captured iModulons represent the known TRN of *Bacillus subtilis* and if the increase in input data resulted in an iModulon decomposition that improves the TRN representation compared to the previous version. Mapping iModulon composition to the known regulon structure of the organism allows quality assessment of the computationally determined iModulon representation. iModulons can be accessed based on how many genes of a specific known regulon were captured in its respective iModulon (recall) and which proportion of the genes that were captured in an iModulon are present in its known regulon counterpart (precision).

Figure 3a shows a bubble plot of recall vs precision of captured iModulons against know regulons and Figure 3b shows a schematic representation of how these values are calculated. The size of each bubble in the plot represents the number of genes in each iModulon. Some interesting iModulons are annotated, like the poorly-matched HrcA iModulon, discussed later. While many of the captured iModulons have high precision, they rarely capturing entire regulons (Regulon-subset). It can be observed that the recall is generally higher for smaller iModulons while large iModulons are often capturing portions of larger regulons like SigB or Rok, which were capture by several iModulons.

**Figure 3:**
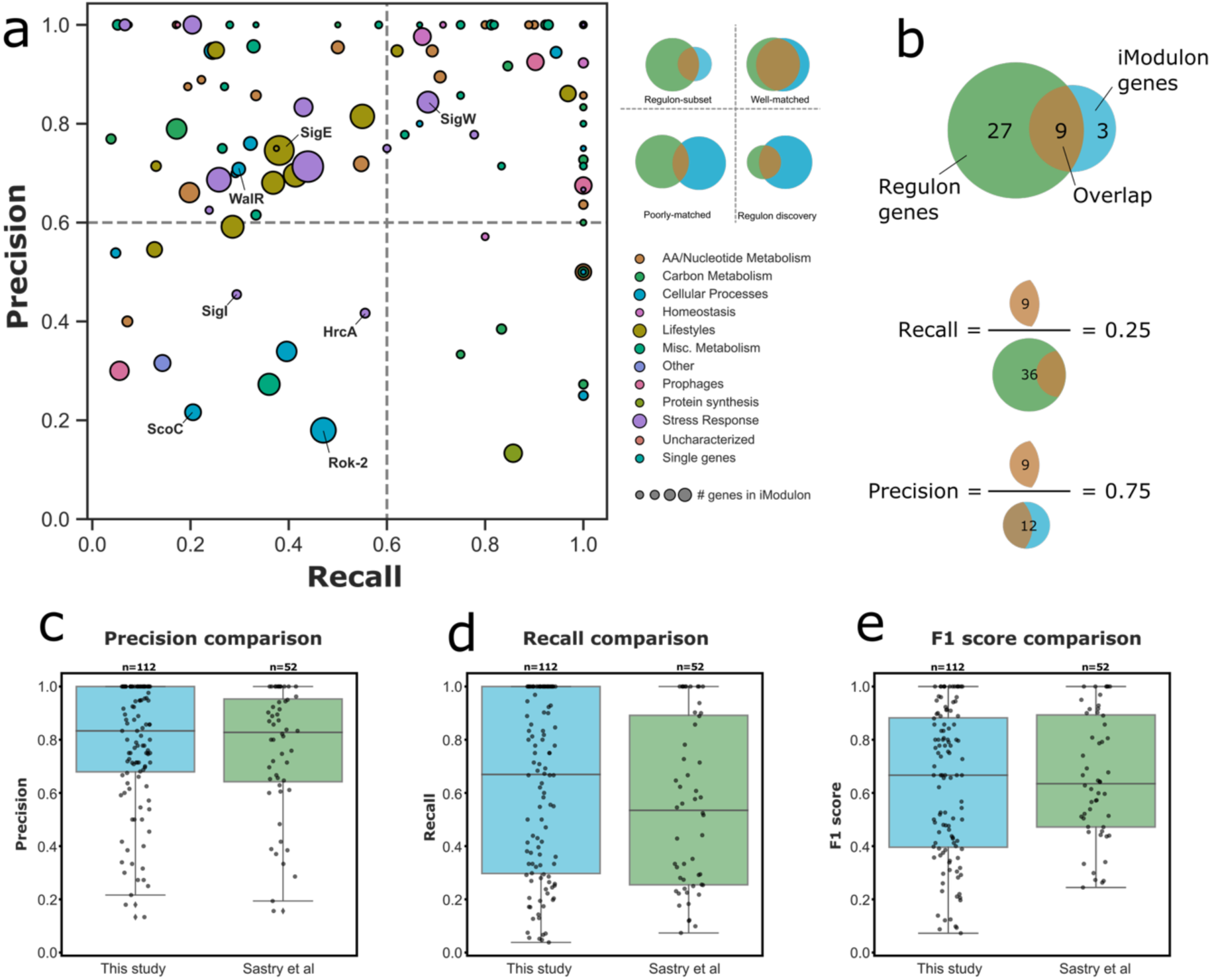
Comparison of iModulons and their regulon counterparts. (a Bubble plot of precision and recall of the captured iModulons in regard to known regulons from *B. subtilis* TRN information. The size of each bubble represents the number of genes in the specific iModulon (iModulon size). The color represents the respective SubtiWiki category the iModulon was associated with. (b) Schematic representation of how recall and precision is calculated from the overlap of captured iModulon and its regulon counterpart. (e-e) Comparison of precision, recall and F1 score the iModulon decomposition presented in this study and the previous iModulon version from Sastry et. al.

Integration of additional public data into the iModulon decomposition resulted in an increase of the median regulon recall from 0.53 to 0.67 (Figure 3d) and consequently in an increase of the median F1 score from 0.63 to 0.67 (Figure 3e), while the iModulon precision stays at 0.83 for both datasets (Figure 3c). The improvement in regulon recall is presumably due to expended condition space and the fact that more regulatory functions including smaller sets of involved genes are captured.

### 4.4 Incorporation of new publicly available RNAseq datasets increases the number of iModulons and the proportion of captured genes and regulons

To investigate the effect of expended data integration in the compositon of iModulons on this new version of the *B. subtilis* TRN decomposition, we compared the updated iModulon version to the version previously reported by Sastry et al. Interestingly, gene members of most previously characterized iModulons were preserved as indicated by Jaccard index of > 0.1 (figure 4a). On the other hand, 53 iModulons and 4 single-gene modules are newly identified in this study, while 2 iModulons and 3 single-gene modules from the previous version are no longer represented. Importantly, iModulons like those representing the regulatory functions of SigB (SigB-1, SigB-2 and SigB-3) were captured in both versions, however, gene contents of these were shuffled. The capturing of 53 novel iModulons can be largely attributed to the presence of more diverse experimental conditions in the input dataset, registering transcriptional changes in previous underrepresented sections of the *B. subtilis* transcriptome. In this updated iModulon decomposition 2383 *Bacillus subtilis* genes were captured, which demonstrates a roughly 20% improvement over the 1981 genes that were captured in iModulons during the study conducted by Sastry et. al (figure 4b). While our updated decomposition captures 142 iModulons, the version of Sastry et al. only captures 72 and the decomposition calculated from microarray date captured 83 iModulons (figure 4c).

**Figure 4:**
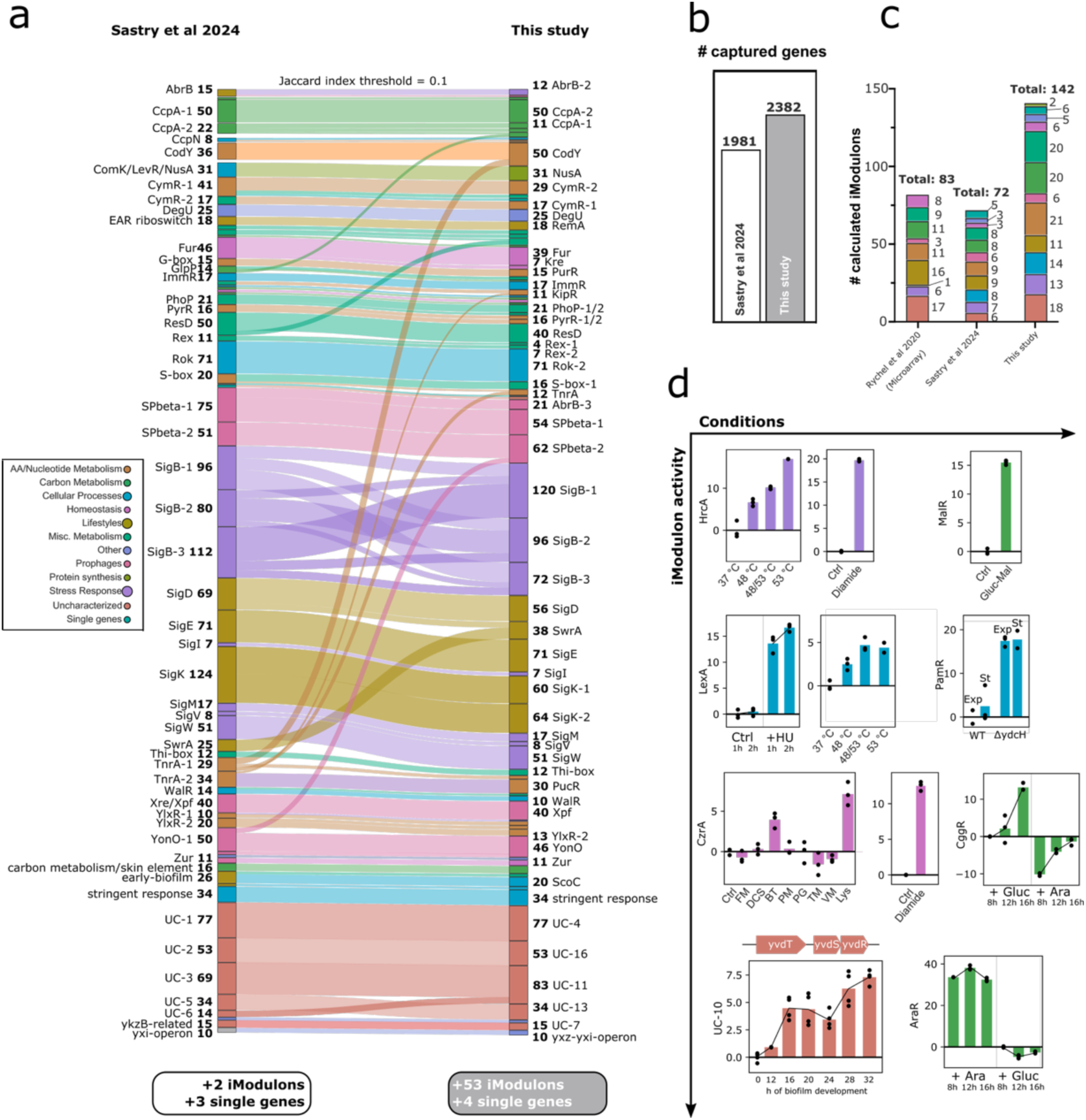
Comparison of iModulon decomposition presented in this study and the previous published version by Sastry et al. (a) Sankey plot of iModulon composition across the iModulons found in both versions of the *B. subtilis* iModulons. The plot shows similarity between the iModulons in both version with a Jaccard index threshold of 0.1, hence representing major rearrangments in gene membership of the respective iModulons. Colors represent respective SubtiWIki categories. (b) Total number of captured genes in both iModulon versions compared in this study. (c) Total number of calculated iModulons in this study in comparison to previous *B. subtilis* iModulon decompositions from Sastry et al. (Sastry et al., 2024) and Rychel et. al (Rychel et al., 2020). Colors represent the respective SubtiWiki categories. (d) iModulon activities of selected novel iModulon captured in this study. Abbreviations in CzrA iModulon activity plot: FM = Flavomycin, DCS = D-cycloserin, BT = Bacitracin, PM = Phosphomycin, PG = Penicillin G, TM = Tunicamycin, VM = Vancomycin, Lys = Lysozyme.

The updated iModulon composition captured several novel iModulons that represent known regulons and previously unknown regulatory functions of known TFs (figure 4d). One example is the HrcA iModulon, which shows increased activity with an increase in cultivation temperature, as well as high activity under diamide treatment conditions. The activities of this novel iModulon corresponds well with known activities and functions of the respective HrcA regulon, which was reported to respond to heat shock conditions and contains chaperons like *groES* and *groEL* (Schumann, 2016). The *hrcA* class I heat stress gene was furthermore reported to respond to disulfide stress induced by diamide treatment, which is also represented in the HrcA iModulon activity levels (Ole Leichert et al., 2003) (figure 4d).

Interestingly the HrcA iModulon showed a poorly-matched recall and precision in comparison to the HrcA regulon. When looking into the gene composition of the HrcA iModulons, it can be noticed that many genes are part of the CtsR regulon. The here reported HrcA iModulon, therefore, captures a mixed signal from HrcA and CtsR regulon (figure S3). The *ctsR* gene was previously reported to encode the transcription repressor of class III heat shock genes like *clpC*, *clpE* and *clpP* and therefore also responds to heat shock conditions (Schumann, 2003). The activation of both regulons under heat shock conditions might explain why the HrcA iModulon capture a mixed signal of two distinct regulons HrcA and CtsR.

Regulatory structures for specific carbon source utilization are captured through novel iModulons. First such iModulon was MalR, showing increased activity upon shift to malate in the cultivation media. Second, CggR iModulon, a known repressor of glycolytic genes, showed a decrease in activity in media supplemented with arabinose instead of glucose as a primary carbon source. Third, AraR iModulon showed increased activity in arabinose cultivation conditions in comparison to glucose as carbon source (Doan et al., 2003), (Bley Folly et al., 2018), (L. Zhang et al., 2012), (Mota et al., 2001). Interestingly, the uncharacterized-10 (UC-10) iModulon a signal comprised of the three operon genes *yvdT*, *yvdS* and *yvdR*. To our knowledge, none of these genes were previously reported to be part of a known regulon. The iModulon activity of UC-10 shows a gradual increase during the development of the *B. subtilis* biofilm (figure 4d). The exact function and role of this iModulon in the developmental process of the organism remain to be elucidated.

### 4.5 iModulons help finding new members of known regulons: WalR regulon

iModulon decomposition of transcriptomics datasets represents a data-driven approach to analyze TRN structures of microorganism. While TRN information, like the one from SubtiWiki, are curated in a bottom-up manner, iModulons enable the scaffolding of such TRNs via computational a top-down approach. Both approaches can complement each other. While known regulon structure can be used to enrich ICA-decomposed signals, novel insights can be drawn from iModulon structures. Consequently, by looking into the gene membership of iModulons in comparison with their regulon counterparts, the here presented computation approach can help identify new members that were previously not known to be regulated by a given TF. We illustrate this by examining the WalR iModulon-regulon relationship.

The WalR iModulon captures a subset equivalent to 28% of the known WalR regulon, which consists of 25 gene-members (figure 5c) (Elfmann et al., 2025), resulting in recall of 0.28 and precision of 0.7. Many of the genes that were not captured by the WalR, were included in other iModulons like the SigI or Rok-2 iModulons, emphasizing how difficult it is to accurately capture the highly interconnected TRN of *Bacillus subtilis*, in which many regulons are substance to crosstalk and genes share membership in multiple regulons. The WalR iModulon captures a set of 10 genes of which 7 are known members of the WalR regulon. Although the WalR iModulon only captures a sub-set of the known WalR regulon, it’s response to different environmental stimuli, corresponds with the known activity of the WalR regulon. As represented in figure 5b, the activity of WalR increases when *B. subtilis* cells are treated with sub-lethal amounts of various cell wall-targeting antibiotics (Q. Zhang et al., 2023). The most notable activity change for WalR iModulon is response to the lysozyme treatment, reaching a 7.6-fold increase over the respective control condition based on the originating work that explores *B. subtilis* cell envelope stress response (Q. Zhang et al., 2023). Additionally, the WalR iModulon shows activity during diamide-induced reactive oxygen species sformation and competency. The iModulons activity strongly aligns with the known activity of the WalR regulon during cell wall homeostasis (Dobihal et al., 2019).

**Figure 5:**
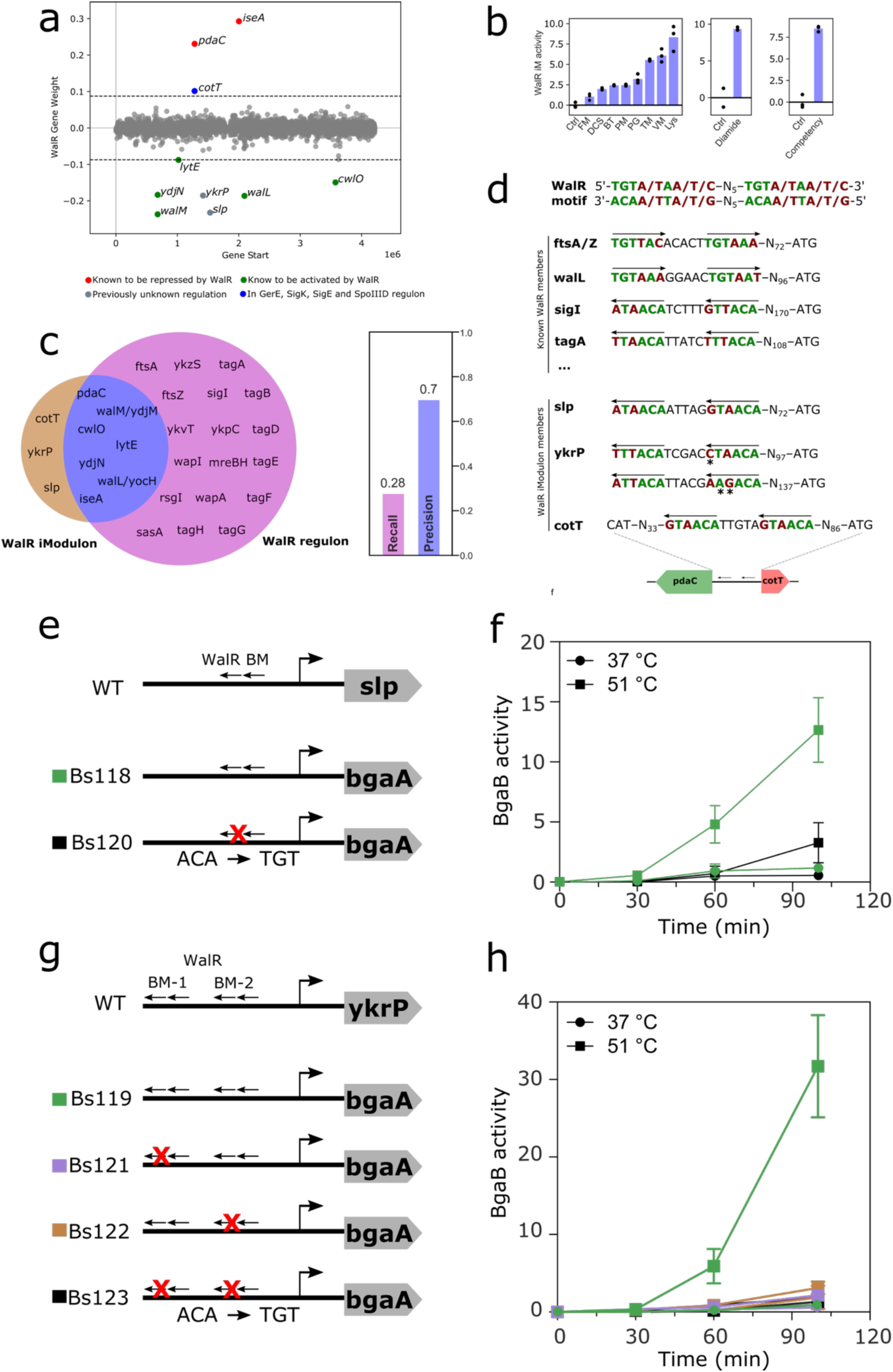
A closer look at the WalR regulon and its gene members. (a) Gene weight plot of genes that were found to be part of the WalR iModulon. Red genes contributing positive gene weight are known to be repressed by WalR. Green genes, showing negative gene weight, are known to be activated by WalR. Grey genes are not associated to any known regulon yet. The blue marked *cotT* gene is known to be regulated the TFs GerE, SigK, SigE and SpoIID. (b) Activity profiles of the WalR iModulon in reponse to different cell wall targeting compounds, diamide and during comptency, FM = Flavomycin, DCS = D-cycloserin, BT = Bacitracin, PM = Phosphomycin, PG = Penicillin G, TM = Tunicamycin, VM = Vancomycin, Lys = Lysozyme (Q. Zhang et al., 2023) (c) Venn diagram showing the overlap of the WalR regulon and the WalR iModulon with *ykrP*, *slp* and *cotT* being previously unknown members of this regulation. (d) Binding motifs of known WalR regulon members and the binding mofis found in upstream of the three genes *slp*, *ykrP* and *cot*. Stars indicated a deviation from the known conserved WalR binding motif. (e and g) Schematic representation of strains cunstructed to investigate the expression mediated by the *slp* and *ykrP* upstream regulayory regions. BM = proposed WalR binding motif. Red crosses indicated the introduction of a mutation within this region. (f and h) BgaB expression regulated by the constructed regulatory native and mutagenic regions in response to cultivation temperature.

As shown by the WalR gene weight scatter plot (figure 5a), the 10 genes contribute to the WalR signal through positive and negative weights. The genes lytE, ydjN, cwlO, walM and walL contribute to the iModulon with a negative weight. Interestingly, all these genes are known to be activate by the WalR regulon (Dobihal et al., 2022), (Huang et al., 2013), (Dobihal et al., 2019). The genes *pdaC* and *iseA* contribute to the iModulon with a positive weight and are known to be repressed by WalR (Dobihal et al., 2022), (Elfmann et al., 2025).

Interestingly, three additional genes are captured by the WalR iModulon, *cotT*, *ykrP* and *slp*, which are not known to be part of the WalR regulon. *cotT,* a gene encoding a protein of the inner spore coat, is known to be a member of several regulons, including GerE, SigK, SigE and SpoIIID. Examining the promoter region of various WalR-regulon members, a binding motif could be found that follows the following structure: 5’-TGTA/TAA/T/C-N_5_-TGTA/TAA/T/C-3’ or 3’-ACAA/T/A/T/G-N_5_-ACAA/TTA/T/G-5’, respectively. This motif was present with various nucleotide distances from the start codon of known WalR regulon members (Howell et al., 2003). Figure 5d depicts some of these known members, including the genes *ftsA/Z, wall*, *sigI* or *tagA*. When looking into the genomic regions of the three potentially new members of this regulon, a similar binding motif was observed, which deviates from the conserved motif by single mutations, only the *slp* upstream region showing a perfectly conserved WalR binding motif. The presence of a WalR binding motif was previously reported via motif search by Howell and co-workers (Howell et al., 2003). It’s transcriptional regulation by WalR, however, is yet to be confirmed. To our knowledge the potential WalR binding motif upstream of *ykrP* was not described previously, and due to nucleotide substitution over the conserved binding motif, most likely not found when performing motif search using the conserved motif sequence. As depicted in figure 5d, there are two potential WalR binding sites upstream of the *ykrP* gene, which vary from the known binding motif by one and two substitutions in the nucleotide sequence respectively. Interestingly, in case of *cotT*, the genes’ location upstream of the known WalR regulon member *pdaC*, might explain its membership in the WalR iModulon, given that’s it’s expression might be indirectly affected through the regulation of its neighboring gene.

To validate the predicted membership of the *slp* and *ykrP* genes in the regulatory network of the WalR TF, we set out to experimentally confirm the binding motif dependent expression of these genes under WalR-relevant growth conditions. As reported previously, WalR is also heat-inducible and fusion of the presumed regulatory regions to the expression of a thermostable β-galactosidase can be used to investigate the expression of genes of interest (Huang et al., 2013). We applied this to the two potential WalR candidates, *slp* and *ykrP*, by amplifying their upstream regulatory regions from *B. subtilis* genomic DNA and cloned them into the pDL backbone, designed to enable genetic fusion of regions of interest to the thermostable β– galactosidase BgaB from *Bacillus stearothermophilus* (Yuan & Wong, 1995). Additionally, mutagenic versions of these plasmids were constructed in which the potential binding motifs (BM) were disrupted. In case of *slp* the native version of the BM version was compared to the mutated version. In case of *ykrP*, the sequence containing two potential BMs (BM-1 and BM-2) was compared to plasmids harboring mutations in either BM-1, BM-2 or bot. A schematic depiction of the constructed *B. subtilis* (168, trpC2 background) strains can be seen in figure 5e and 5g. As shown in figure 5f and 5h, the expression of the *bgaB gene* fused to the *slp* and *ykrP* upstream binding motifs is activated once the strain is cultivated under heat conditions (51 °C) and absence when cultivated in 37 °C. This activation is alleviated once the presumed binding motifs are mutated (single and in combination for *ykrP*). Due to the heat-inducible expression of *slp* and *ykrP* which is alleviated of expression upon mutation of the WalR-BM resembeling upstream regions, this result suggests that both s*lp* and *ykrP* might be activated by WalR as part of the cell wall homeostasis regulation.

### 5.5 A model describing the potential role of ykrP in the WalR-mediated cell wall homeostasis cycle

Based on the membership of *slp* and *<u>ykrP</u>* in the WalR iModulon and their WalR-binding-motif-dependent expression under high-temperature condition, which was previously shown to induce the expression of the *walR* gene and WalR regulon (Huang et al., 2013), we propose new hypothesis regarding the roles of these genes (Figure 6).

**Figure 6:**
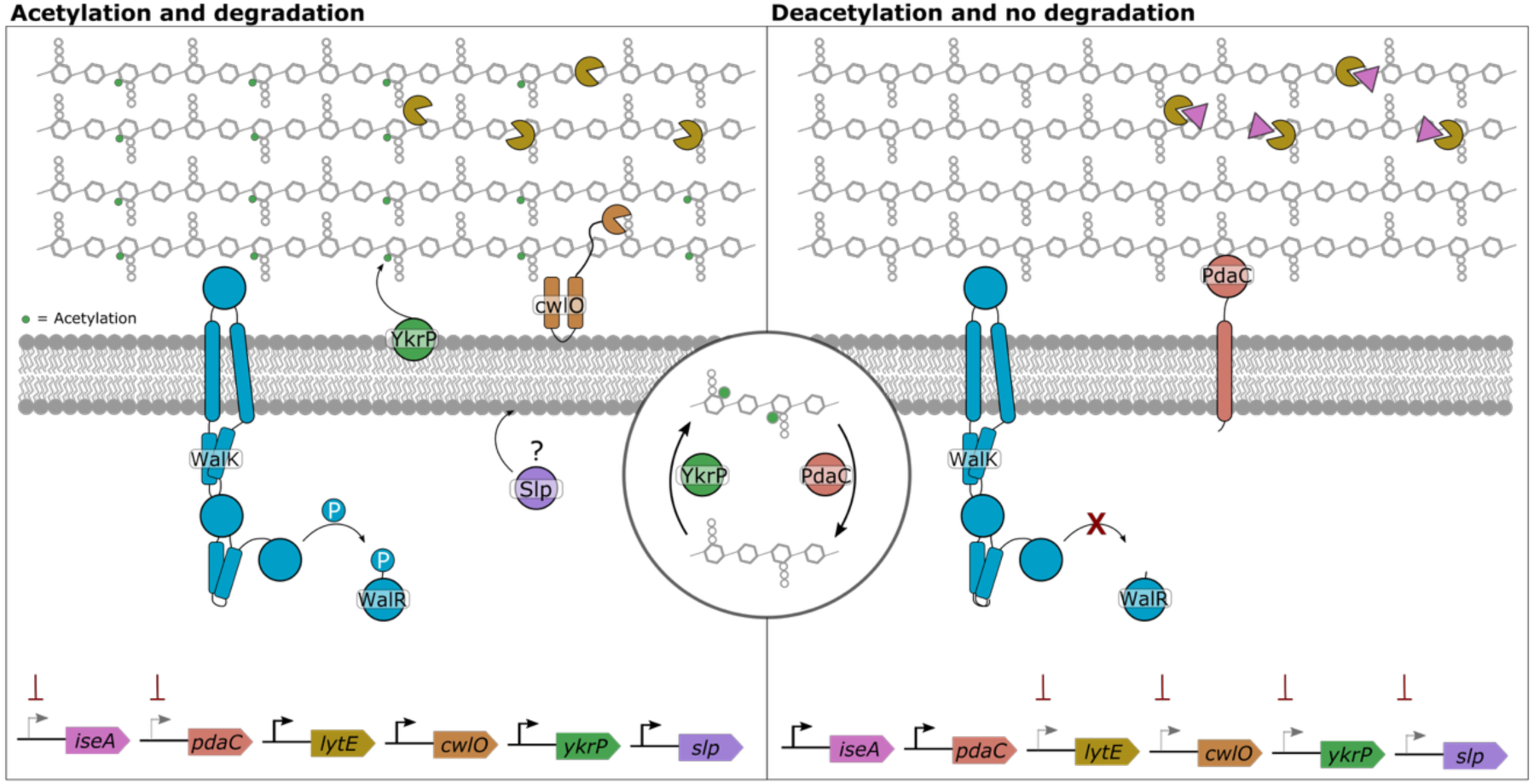
Schematic model describing the potential regulatory role of WalR on the expression of ykrP and slp with the ykrp gene product YkrP acting as a peptidoglycan acetylase competing with the known peptidoglycan deactylase PdaC in a cycle of acetylation and deacylation, controlling the level of peptidogylcan degradation during cell wall homeostasis in *Bacillus subtilis*. The figure was adapted and extended from (Dobihal et al., 2022).

The *ykrP*-encoded membrane-bound acyltransferase YkrP is predicted to contain an acetyltransferase 3 (AT-3) domain, which is known to be present in enzymes that catalize the transfer of acyl group, e.g. OatA and YfiQ in B. subtilis. Notably, OatA a O-acetyl transferase that provides resistance to lysozyme, catalyze the transfer of acyl groups with OatA acting on peptidoglycan in the well membrane (Veronica & D, 2011), (Jones et al., 2020).

The WalR TF is known to be able to regulate gene expression both positively through activation and negatively via repression. WalR is known to repress the peptidoglycan deacetylase PdaC (Dobihal et al., 2022). The negative gene weight contributed by *ykrP* to the WalR iModulon suggests that this gene, among the other known activated genes, is also activated by WalR. Based on these findings we propose a counteracting function of YkrP reversing the diacylation performed by PdaC and therefore contributing to cell wall homeostasis and the WalR-mediated response to environmental stimuli like heat, lysozyme treatment or cell wall targeting antibiotics (figure 6). The 375 bp *slp* gene is predicted to encode a small peptidoglycan-associated lipoprotein with 124 amino acids in size. The potential function of this gene is yet to be elucidated. Further analysis needs to be conducted to link the expression of both *ykrP* and *slp* to the WalR regulon and might be confirmed via electromobility shift assays or CHIPseq.

## 5. Conclusion

In this study, we presented an updated iModulon decomposition for the transcriptome of the model organism and industrial workhorse for protein production, *Bacillus subtilis*, leveraging an expanded compendium of 782 high-quality RNAseq samples, which were mined from the publicly available NCBI SRA database. By applying independent component analysis to this compendium, we could identify 142 independent regulated sets of genes – iModulons, which explain 80% of the expression variance in the underlying input dataset. The decomposition manages to capture many known regulators robustly and the activity of the iModulons accurately reflects the known mode of action of the associated regulons counterparts. The increase from previously 72 iModulons captured by Sastry et. al to 142 robust iModulons not only captures a broader array of environmental and genetic perturbations but also improves the resolution of known regulatory modules, revealing biologically meaningful gene regulations. Our study described 53 novel iModulons, previous not captured in ICA decompositions, many of which are active under previously underrepresented experimental conditions. With the newly captured iModulon decomposition at hand, we investigated its utilization for exploring previously unknown regulatory relationships.

Our study has also demonstrated this approach can provide novel insights into well-characterized regulons. The WalR iModulon captured genes-*slp* and *ykrP*, previously not recognized as part of the WalR regulon. Closer inspection of the upstream regulatory regions of these genes revealed potential WalR DNA binding motifs. Experimental investigated showed that mutagenesis of these binding motifs, alleviates the activation of the downstream genes upon heat shock, a condition known to be activate the WalR regulon response. These results supported the regulation of *slp* and *ykrP* genes WalR regulator. We further proposed a model of cell-wall homeostasis incorporating the *ykrP* gene.

As the volume of transcriptomics data continues to grow, this study underscores the importance of maintaining and updating community-curated resources such as iModulonDB. Further integration of complementary data types such as proteomics, metabolomics and CHIPseq could further enrich the iModulon framework and deepen our understanding of transcriptional regulation in *B. subtilis* and related Gram-positive bacteria.

## Author Contributions

P.G. wrote the original draft of the article, produced visualizations and conducted data analysis and experimental work. P.G., E. Ö. and L. Y. conceived the study. P. G., E. Ö., L.Y. and D. Z. analyzed and interpreted results. E. Ö., L.Y., I.M., B.P., supervised and managed the project. All authors have reviewed and agreed to the published version of the article.

## Acknowledgments

This work was funded by the Novo Nordisk Foundation grant NNF20CC0035580.

## References

Bley Folly, B., Ortega, A. D., Hubmann, G., Bonsing-Vedelaar, S., Wijma, H. J., van der Meulen, P., Milias-Argeitis, A., & Heinemann, M. (2018). Assessment of the interaction between the flux-signaling metabolite fructose-1,6-bisphosphate and the bacterial transcription factors CggR and Cra. Molecular Microbiology, 109(3), 278–290. 10.1111/mmi.14008

Catoiu, E. A., Krishnan, J., Li, G., Lou, X. A., Rychel, K., Yuan, Y., Bajpe, H., Patel, A., Choe, D., Shin, J., Burrows, J., Phaneuf, P. V., Zielinski, D. C., & Palsson, B. O. (2025). IModulonDB 2.0: Dynamic tools to facilitate knowledge-mining and user-enabled analyses of curated transcriptomic datasets. Nucleic Acids Research, 53(D1), D99–D106. 10.1093/nar/gkae1009

Doan, T., Servant, P., Tojo, S., Yamaguchi, H., Lerondel, G., Yoshida, K. I., Fujita, Y., & Aymerich, S. (2003). The Bacillus subtilis ywkA gene encodes a malic enzyme and its transcription is activated by the YufL/YufM two-component system in response to malate. Microbiology, 149(9), 2331–2343. 10.1099/mic.0.26256-0

Dobihal, G. S., Brunet, Y. R., Flores-Kim, J., & Rudner, D. Z. (2019). Homeostatic control of cell wall hydrolysis by the WalRK two-component signaling pathway in Bacillus subtilis. ELife, 8. 10.7554/eLife.52088

Dobihal, G. S., Flores-Kim, J., Roney, I. J., Wang, X., & Rudner, D. Z. (2022). The WalR- WalK Signaling Pathway Modulates the Activities of both CwlO and LytE through Control of the Peptidoglycan Deacetylase PdaC in Bacillus subtilis.

Elfmann, C., Dumann, V., van den Berg, T., & Stülke, J. (2025). A new framework for SubtiWiki, the database for the model organism Bacillus subtilis. Nucleic Acids Research, 53(D1), D864–D870. 10.1093/nar/gkae957

Ewels, P., Magnusson, M., Lundin, S., & Käller, M. (2016). MultiQC: summarize analysis results for multiple tools and samples in a single report. Bioinformatics, 32(19), 3047–3048. 10.1093/bioinformatics/btw354

Gu, Y., Xu, X., Wu, Y., Niu, T., Liu, Y., Li, J., Du, G., & Liu, L. (2018). Advances and prospects of Bacillus subtilis cellular factories: From rational design to industrial applications. In Metabolic Engineering (Vol. 50, pp. 109–121). Academic Press Inc. 10.1016/j.ymben.2018.05.006

Harwood, C. R. (1992). Bacillus subtilis and its relatives: molecular biological and industrial workhorses. Trends in Biotechnology, 10, 247–256. 10.1016/0167-7799(92)90233-L

Howell, A., Dubrac, S., Andersen, K. K., Noone, D., Fert, J., Msadek, T., & Devine, K. (2003). Genes controlled by the essential YycG/YycF two-component system of Bacillus subtilis revealed through a novel hybrid regulator approach. Molecular Microbiology, 49(6), 1639–1655. 10.1046/j.1365-2958.2003.03661.x

Huang, W.-Z., Wang, J.-J., Chen, H.-J., Chen, J.-T., & Shaw, G.-C. (2013). The heat-inducible essential response regulator WalR positively regulates transcription of sigI, mreBH and lytE in Bacillus subtilis under heat stress. Research in Microbiology, 164(10), 998–1008. 10.1016/j.resmic.2013.10.003

Hyvarinen, A. (1999). Fast ICA for noisy data using Gaussian moments. 1999 IEEE International Symposium on Circuits and Systems (ISCAS), 5, 57–61 vol.5. 10.1109/ISCAS.1999.777510

Jones, C. S., Sychantha, D., Howell, P. L., & Clarke, A. J. (2020). Structural basis for the *O*-acetyltransferase function of the extracytoplasmic domain of OatA from *Staphylococcus aureus*. Journal of Biological Chemistry, 295(24), 8204–8213. 10.1074/jbc.RA120.013108

Jönsson, M., Sigrist, R., Gren, T., Semenov Petrov, M., Marcussen, N. E. J., Svetlova, A., Charusanti, P., Gockel, P., Palsson, B. O., Yang, L., & Özdemir, E. (2025). Machine learning uncovers the transcriptional regulatory network for the production host Streptomyces albidoflavus. Cell Reports, 44(3). 10.1016/j.celrep.2025.115392

Katz, K., Shutov, O., Lapoint, R., Kimelman, M., Rodney Brister, J., & O’Sullivan, C. (2022). The Sequence Read Archive: A decade more of explosive growth. Nucleic Acids Research, 50(D1), D387–D390. 10.1093/nar/gkab1053

Langmead, B., & Salzberg, S. L. (2012). Fast gapped-read alignment with Bowtie 2. Nature Methods, 9(4), 357–359. 10.1038/nmeth.1923

Liao, Y., Smyth, G. K., & Shi, W. (2014). featureCounts: an efficient general purpose program for assigning sequence reads to genomic features. Bioinformatics, 30(7), 923–930. 10.1093/bioinformatics/btt656

Liu, Z.-Y., & Yu, X.-Z. (2025). Engineering Bacillus subtilis for high-value bioproduction: recent advances and applications. Microbial Cell Factories, 24(1), 182. 10.1186/s12934-025-02818-6

McConn, J. L., Lamoureux, C. R., Poudel, S., Palsson, B. O., & Sastry, A. V. (2021). Optimal dimensionality selection for independent component analysis of transcriptomic data. BMC Bioinformatics, 22(1), 584. 10.1186/s12859-021-04497-7

Mota, L. J., Morais Sarmento, L., & De Sá-Nogueira, I. (2001). Control of the arabinose regulon in Bacillus subtilis by AraR in vivo: Crucial roles of operators, cooperativity, and DNA looping. Journal of Bacteriology, 183(14), 4190–4201. 10.1128/JB.183.14.4190-4201.2001

Ole Leichert, L. I., Scharf, C., & Hecker, M. (2003). Global characterization of disulfide stress in Bacillus subtilis. Journal of Bacteriology, 185(6), 1967–1975. 10.1128/JB.185.6.1967-1975.2003

Rychel, K., Sastry, A. V., & Palsson, B. O. (2020). Machine learning uncovers independently regulated modules in the Bacillus subtilis transcriptome. Nature Communications, 11(1). 10.1038/s41467-020-20153-9

Saint-André, V. (2021). Computational biology approaches for mapping transcriptional regulatory networks. In Computational and Structural Biotechnology Journal (Vol. 19, pp. 4884–4895). Elsevier B.V. 10.1016/j.csbj.2021.08.028

Sastry, A. V., Gao, Y., Szubin, R., Hefner, Y., Xu, S., Kim, D., Choudhary, K. S., Yang, L., King, Z. A., & Palsson, B. O. (2019). The Escherichia coli transcriptome mostly consists of independently regulated modules. Nature Communications, 10(1). 10.1038/s41467-019-13483-w

Sastry, A. V., Yuan, Y., Poudel, S., Rychel, K., Yoo, R., Lamoureux, C. R., Li, G., Burrows, J. T., Chauhan, S., Haiman, Z. B., Bulushi, T. Al, Seif, Y., Palsson, B. O., & Zielinski, D. C. (2024). iModulonMiner and PyModulon: Software for unsupervised mining of gene expression compendia. PLoS Computational Biology, 20(10 October). 10.1371/journal.pcbi.1012546

Schumann, W. (2003). The Bacillus subtilis heat shock stimulon. In Cell Stress & Chaperones (Vol. 8, Issue 3). Cell Stress Society International.

Schumann, W. (2016). Regulation of bacterial heat shock stimulons. Cell Stress and Chaperones, 21(6), 959–968. 10.1007/s12192-016-0727-z

Shin, J., Rychel, K., & Palsson, B. O. (2023). Systems biology of competency in Vibrio natriegens is revealed by applying novel data analytics to the transcriptome. Cell Reports, 42(6). 10.1016/j.celrep.2023.112619

Stülke, J., Grüppen, A., Bramkamp, M., & Pelzer, S. (2023). Bacillus subtilis, a Swiss Army Knife in Science and Biotechnology. In Journal of Bacteriology (Vol. 205, Issue 5). American Society for Microbiology. 10.1128/jb.00102-23

Veronica, G.-O., & D, H. J. (2011). Bacillus subtilis σV Confers Lysozyme Resistance by Activation of Two Cell Wall Modification Pathways, Peptidoglycan O-Acetylation and d-Alanylation of Teichoic Acids. Journal of Bacteriology, 193(22), 6223–6232. 10.1128/jb.06023-11

Wang, L., Wang, S., & Li, W. (2012). RSeQC: quality control of RNA-seq experiments. Bioinformatics, 28(16), 2184–2185. 10.1093/bioinformatics/bts356

Westers, L., Westers, H., & Quax, W. J. (2004). Bacillus subtilis as cell factory for pharmaceutical proteins: A biotechnological approach to optimize the host organism. In Biochimica et Biophysica Acta - Molecular Cell Research (Vol. 1694, Issues 1-3 SPEC.ISS., pp. 299–310). 10.1016/j.bbamcr.2004.02.011

Yuan, G., & Wong, S. L. (1995). Regulation of groE expression in Bacillus subtilis: the involvement of the sigma A-like promoter and the roles of the inverted repeat sequence (CIRCE). Journal of Bacteriology, 177(19), 5427–5433. 10.1128/jb.177.19.5427-5433.1995

Zhang, L., Leyn, S. A., Gu, Y., Jiang, W., Rodionov, D. A., & Yang, C. (2012). Ribulokinase and transcriptional regulation of arabinose metabolism in clostridium acetobutylicum. Journal of Bacteriology, 194(5), 1055–1064. 10.1128/JB.06241-11

Zhang, Q., Cornilleau, C., Müller, R. R., Meier, D., Flores, P., Guérin, C., Wolf, D., Fromion, V., Carballido-Lopez, R., & Mascher, T. (2023). Comprehensive and Comparative Transcriptional Profiling of the Cell Wall Stress Response in *Bacillus subtilis* BioRxiv, 2023.02.03.526509. 10.1101/2023.02.03.526509

