## Supplementary Figures for "Expanding the iModulon Knowledgebase for *Bacillus subtilis*: Updated Transcriptome Decomposition Reveals Novel Regulatory Mechanisms"

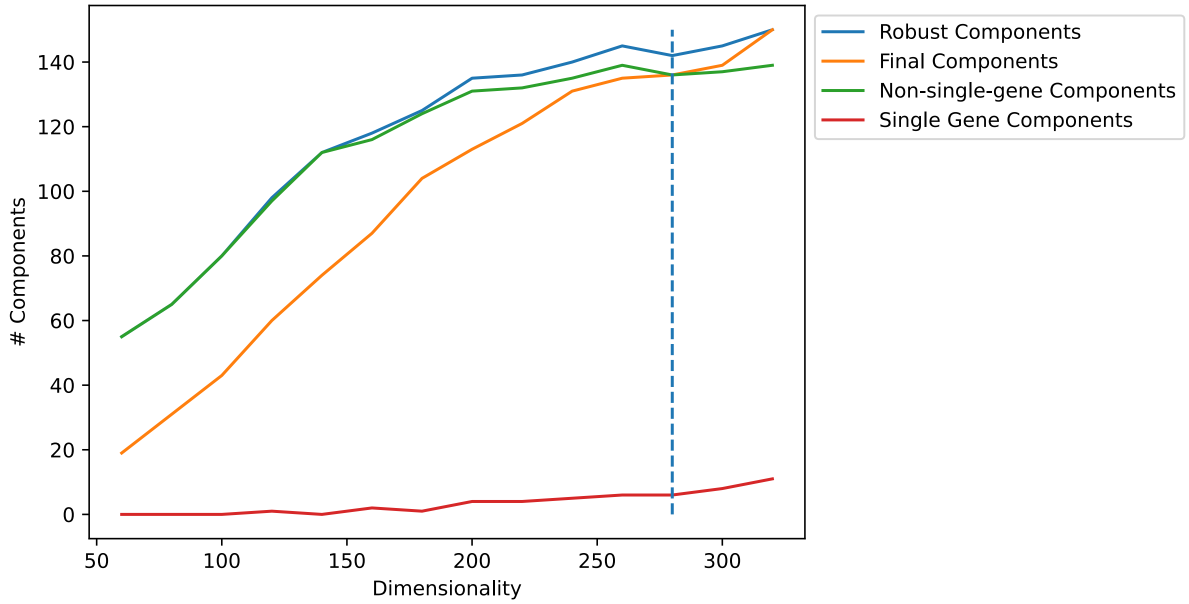


**Figure S1:** Dimension analysis during optICA process. The optimal number of dimensions was found to be 280 resulting in 142 robust components.

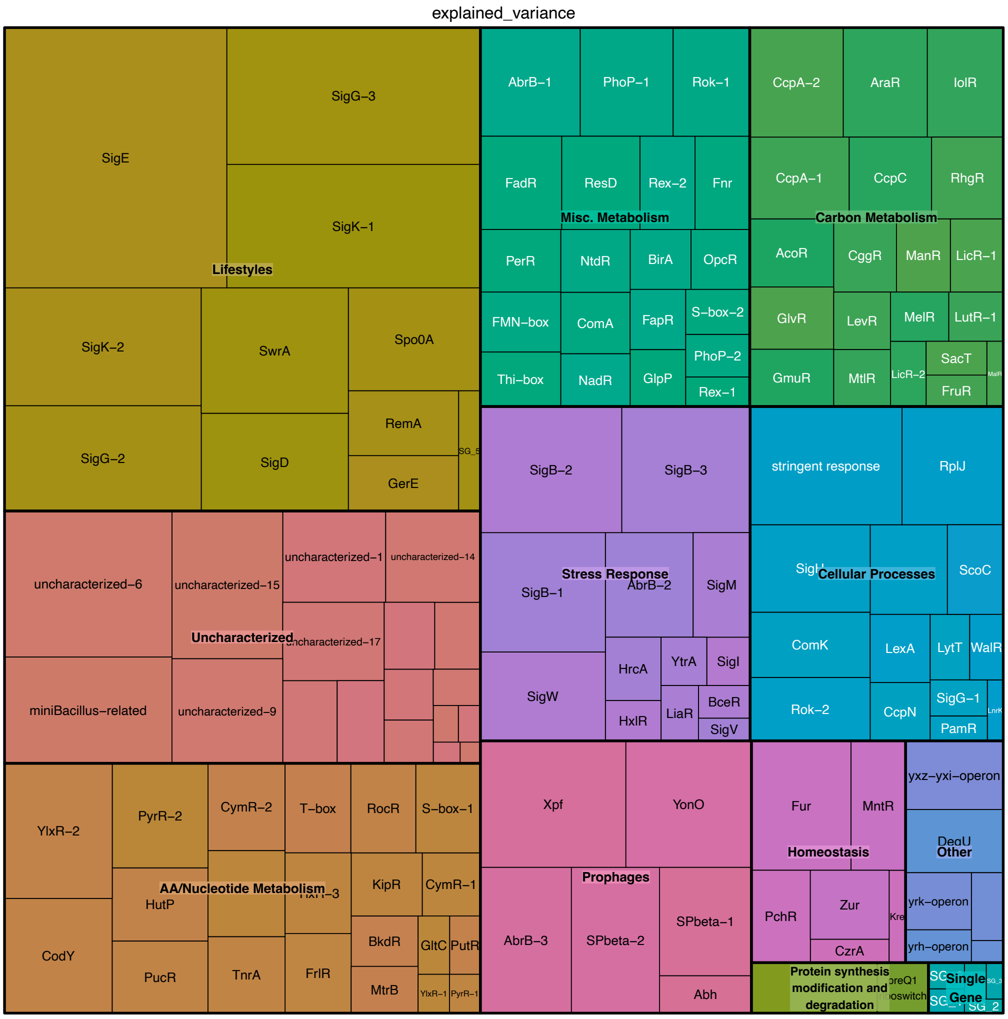


**Figure S2:** Treemap of iModulons in which each square represent the explained variance contributed from the respective iModulon.

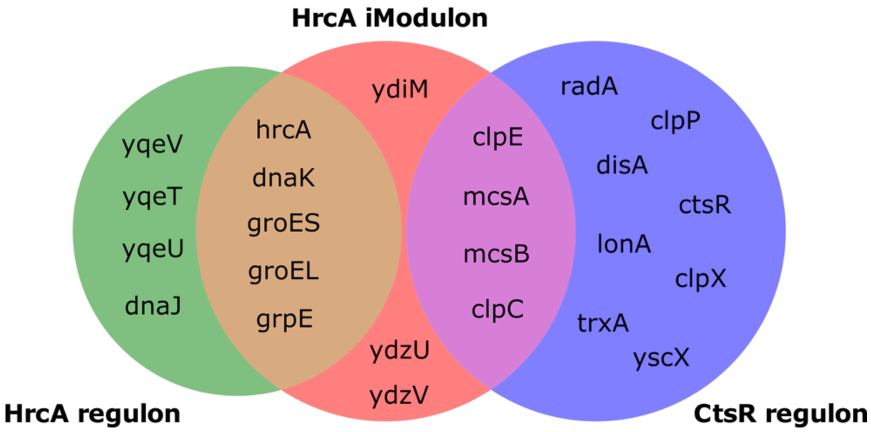


**Figure S3:** Venn diagram of HrcA iModulon and the respective known HrcA and CtsR regulon.
